# Plant DNA Designer: A Computational Framework for Multi-Objective Codon Optimisation and Synthetic Gene Design in Crop Biotechnology

**DOI:** 10.64898/2026.08.02.742271

**Authors:** K Dinesh, H. Swetha

## Abstract

Synthetic gene design for plant transformation requires simultaneous optimisation of multiple, often competing, molecular objectives: translational efficiency, mRNA structural accessibility, codon-pair compatibility, regulatory safety, and species-specific expression context. Existing tools address these objectives in isolation, typically maximising a single metric such as the Codon Adaptation Index (CAI) and neglecting the broader determinants of in-plant expression. We present **Plant DNA Designer (PDD)**, a web-based platform that integrates a 19-objective genetic algorithm with expression-cassette co-design, clade-aware translation-initiation logic, ribosome-velocity trajectory shaping, CRISPR guide-RNA design, and multi-gene pathway balancing across 18 crop species spanning monocot and dicot clades — each using its own measured codon-usage table from the Kazusa Codon Usage Database. We benchmark PDD against faithful reproductions of the published algorithms of five external tools (JCat/OPTIMIZER/ATGme, IDT, TISIGNER, a CAI+GC heuristic, and a random floor) across six validated rice effector proteins. PDD is the only strategy that holds every objective within acceptable bounds at once: it reduces transgene safety liabilities from 2.3–3.5 to 0.0, and cuts deviation from a 50 % GC synthesis target from 21.8 to 4.0 percentage points, while raising codon harmony from 0.42 to 0.77 — at a deliberate, moderate cost in raw CAI (0.79 vs 1.00). Consistent with a fair comparison rather than a strawman, a dedicated single-objective tool (IDT) still outperforms PDD on its own axis (harmony 0.93). We anchor the two central proxies against real biology: on 456 real rice genes, CAI and the wobble-weighted tAI are significantly higher in highly-expressed ribosomal-protein genes than in the genomic background (Mann–Whitney *p* ≤ 10⁻⁵; tAI AUC 0.75) and correlate at Spearman *ρ* = 0.93. Beyond this expression-class anchor, the reported design metrics are in-silico proxies, not wet-lab yield measurements. PDD is released as open-source software under an MIT licence and is freely accessible as a FastAPI web application.

## Paper at a glance

| Section | What you will find |
| --- | --- |
| 1. Introduction | What existing codon tools do, the gap they leave, and how PDD closes it |
| 2. Background and Related Work | The biology and prior methods: CAI/tAI/harmony, mRNA structure, plant Kozak/IME, cassettes, CRISPR, pathways |
| 3. The Plant DNA Designer System | The methods — architecture, the 19-objective fitness function, the genetic algorithm, plant-specific logic, cassette and pathway modules, and (3.9) software versions, parameters and reproducibility |
| 4. Results | Benchmark against five external-tool algorithms (with significance); empirical validation of CAI/tAI on real rice genes; species-specific, GA/Pareto, and pathway results |
| 5. Discussion | Why multi-objective beats single-metric design, plant-specific considerations, and limitations (incl. no wet-lab validation) |
| 6. Conclusion | Summary of contributions and outcomes |

## 1. Introduction

Plant synthetic biology has matured from proof-of-concept demonstrations into a central engineering discipline for crop improvement (Patron et al., 2015). The ability to design, synthesise, and deliver heterologous DNA constructs that express predictably in a chosen crop species underpins a broad range of agricultural applications: pathogen resistance, abiotic-stress tolerance, nutrient biofortification, yield enhancement, and bioenergy optimisation (Ye et al., 2000; Halpin, 2005). Yet despite decades of progress, the gap between computational DNA design and reliable in-planta expression remains large. Constructs that appear optimal by standard metrics routinely underperform in transformation experiments, and the failure modes are poorly understood by practitioners who rely on single-metric tools.

The fundamental challenge is that the genetic code is degenerate: most amino acids can be encoded by two to six synonymous codons, creating an enormous synonymous sequence space (4ⁿ for a sequence of *n* codons) that must be navigated to maximise expression. The Codon Adaptation Index (CAI; Sharp and Li, 1987), computed as the geometric mean of per-codon relative adaptiveness values with respect to a reference set of highly expressed genes, has long served as the standard proxy for translational fitness. However, as recent systematic comparisons demonstrate, tools that maximise CAI in isolation fail to account for mRNA secondary structure stability, codon-pair bias, tRNA availability, and the structured initiation window at the 5′ end of the transcript (Kudla et al., 2009; Tuller et al., 2010; Mauro and Chappell, 2014). The result is flat-fast sequences that saturate tRNA pools, lose the natural ramp at the start of translation, and accumulate forbidden restriction sites and forbidden motifs that limit downstream cloning.

Plant systems introduce additional complexity not seen in microbial expression. The plant ribosome operates under a distinct Kozak context that differs between monocotyledonous and dicotyledonous plants (Joshi et al., 1987; Sawant et al., 2001). Intron-mediated enhancement (IME) — the increase in mRNA accumulation and translational efficiency conferred by a suitably positioned intron — is a plant-specific phenomenon with clade-dependent sequence requirements (Mascarenhas et al., 1990; Chung et al., 2006). Plant miRNAs targeting transgene sequences can silence transgene expression post-transcriptionally (Palatnik et al., 2003). Expression cassettes must be assembled from validated regulatory parts — promoters, 5′ leaders, and terminators — whose function has been confirmed experimentally; *de-novo* regulatory elements remain unreliable (Odell et al., 1985; Gallie et al., 1987). Finally, multi-gene metabolic pathway engineering requires balanced, coordinated expression across several coding sequences, which single-gene tools cannot address.

### What existing tools do

General-purpose codon optimisers — JCat (Grote et al., 2005), OPTIMIZER (Puigbò et al., 2007), ATGme, the commercial IDT and GeneArt optimisers, and TISIGNER (Bhandari et al., 2021) — each optimise one dimension of the problem: most maximise CAI by choosing the single most-frequent codon per residue, IDT balances codon frequencies, and TISIGNER tunes the translation-initiation window. They are predominantly built for microbial or mammalian hosts and treat the coding sequence in isolation from its regulatory cassette.

### What gap remains

No existing tool simultaneously (i) optimises the competing molecular determinants of expression — codon adaptation, codon harmony, tRNA adaptation, position-dependent mRNA structure, ribosome-velocity dynamics, and cloning safety — under one objective; (ii) applies *plant-specific* biology end-to-end (monocot/dicot Kozak context, clade-appropriate IME introns, plant miRNA avoidance, and per-species codon usage); or (iii) extends from the coding sequence to the full validated cassette and to balanced multi-gene pathways. Practitioners therefore stitch together several single-purpose, host-generic tools and still produce sequences that mis-express in planta.

### Why Plant DNA Designer addresses it

PDD closes this gap by treating plant gene design as a single multi-objective optimisation problem, solved by a genetic algorithm whose 19 fitness components encode exactly these competing determinants, with plant- and clade-specific biology applied throughout and the design extended to the full cassette and to multi-gene pathways.

Here we describe Plant DNA Designer (PDD), a comprehensive computational platform that addresses all of these requirements in a single, integrated web application. PDD implements a multi-objective genetic algorithm with 19 fitness components, each normalised to a dimensionless score and weighted according to biological importance. The platform includes: (i) a 19-objective fitness function that replaces single-metric CAI maximisation; (ii) a position-dependent mRNA structural model that distinguishes beneficial body stability from harmful 5′-end structure; (iii) a translation-dynamics model that explicitly shapes the ribosome velocity trajectory; (iv) species-specific expression logic covering 18 crops across monocot and dicot clades; (v) a full expression cassette co-design module with validated part libraries; (vi) a multi-gene pathway balancing module using codon-optimality dialing; and (vii) CRISPR/Cas guide RNA design for all 18 supported crop species.

The remainder of the paper is organised as follows. Section 2 reviews the relevant literature on codon optimisation, translation initiation, and plant expression engineering. Section 3 describes the PDD system in detail. Section 4 presents benchmark results. Section 5 discusses the implications and limitations. Section 6 concludes.

## 2. Background and Related Work

### 2.1 Codon Optimisation Metrics

The CAI (Sharp and Li, 1987) quantifies how similar a coding sequence is to the codon usage pattern of a reference set — typically the most highly expressed genes in the target organism. Given a set of synonymous codons {*c*₁,…, *c*ₙ} for amino acid *AA*, the relative adaptiveness of codon *c* is w(*c*) = f(*c*) / max{f(*c*ⱼ)}, where f(*c*) is the frequency of codon *c* among synonymous alternatives in the reference. CAI for a coding sequence of *L* codons is:

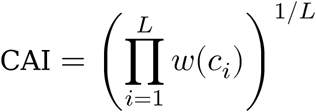

Systematic benchmarking of ten codon optimisation tools found that tools maximising CAI (JCat, OPTIMIZER, ATGme) consistently achieved CAI ≈ 1.0 but failed to account for mRNA secondary structure stability and codon-pair bias, which profoundly influence protein expression, especially for longer sequences (Mauro and Chappell, 2014). These observations confirm the inadequacy of a single-metric approach and motivate multi-parameter strategies.

The tRNA Adaptation Index (tAI; dos Reis et al., 2004) addresses a fundamental limitation of CAI: it uses codon-anticodon interaction efficiency weights derived from tRNA gene copy numbers (tGCN) in the target genome, rather than frequency in highly expressed genes. For each codon *c* decoded by anticodon *a*, the absolute adaptiveness is:

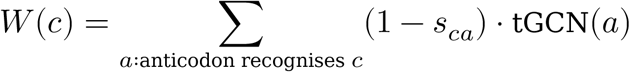

where *s*_{ca} is the codon–anticodon wobble penalty (dos Reis et al. 2004 optimised values: I:C = 0.28, I:A = 0.9999, G:U = 0.41, U:G = 0.68; Watson–Crick pairings = 0), so a codon read only through an inefficient wobble tRNA is down-weighted relative to its Watson–Crick synonym. The per-codon relative adaptiveness *w(c) = W(c)/max_c W(c)* is precomputed per species (`tools/build_tgcn.py`); codons with no decoding tRNA take the geometric-mean fill of the non-zero weights, so a tRNA-less codon lowers rather than escapes the score. The tAI geometric mean over all codons gives a score that correlates with empirical protein abundance independently of CAI. PDD uses per-species tGCN sets counted directly from GtRNAdb genome annotations (anticodon gene counts expanded to codons under standard eukaryotic wobble with A34→I34 inosine): 11 crops have their own sequenced genome, and the remaining crops map to their nearest sequenced relative by phylogeny (e.g. potato→tomato, peanut→soybean, sugarcane→sorghum, wheat and barley→*Brachypodium distachyon*). The source FASTAs are committed and the tables are regenerated by `tools/build_tgcn.py`.

#### Codon Harmony

(Angov, 2011; Mignon et al., 2018) inverts the CAI logic: instead of maximising use of the most-frequent codon, it measures the total-variation distance between the design’s per-amino-acid codon-usage distribution and the host reference distribution. A design with identical distribution to the reference scores 1.0 (perfect harmony); a design using only the most-frequent codons for each amino acid scores ∼0.4. The rationale is that natural synonymous codon usage encodes secondary information — tRNA trafficking, ribosome pausing for co-translational folding — that CAI-maximisation destroys.

### 2.2 mRNA Secondary Structure

Early work assumed that mRNA secondary structure was uniformly deleterious, but the picture is more nuanced. Kudla et al. (2009) demonstrated that 5′ UTR and start-codon region structure (roughly the first 45 nt of the CDS) strongly predicts protein expression level across a library of 154 green fluorescent protein variants in *E. coli*; sequences with low Minimum Free Energy (MFE) in this window expressed at higher levels. Conversely, moderate secondary structure in the body of the transcript is not deleterious and can protect the mRNA from degradation, so structure must be evaluated position-by-position rather than globally minimised (Mauro and Chappell, 2014).

These results define two distinct structural objectives that a complete optimiser must handle: **start accessibility** (the 5′ first ∼45 nt must be structurally open, ΔG → 0 or positive) and **body stability** (the remainder of the mRNA benefits from modest secondary structure, ΔG more negative). PDD implements both as separate windowed sub-scores.

### 2.3 Translation Initiation Context in Plants

The Kozak consensus — the nucleotide context surrounding the AUG start codon — modulates cap-dependent translation initiation efficiency. The optimal context differs between plant clades. Joshi et al. (1987, 1997) analysed plant mRNA sequences and identified an A-rich upstream context with the consensus A/CAAUGGC, where the −3 position adenine is most critical. Sawant et al. (2001) conducted the analogous analysis for monocots (rice, maize, wheat) and found a GC-richer context, reflecting the higher overall GC content of grass genomes. These clade-specific Kozak tables — incorporated in PDD — can shift predicted translation initiation efficiency by 15–25% relative to a generic (mammalian) Kozak consensus.

Intron-Mediated Enhancement (IME) is a plant-specific phenomenon in which introns positioned in the 5′ leader or first exon increase expression 2–10-fold (Mascarenhas et al., 1990). The IMEter scoring model (Rose et al., 2008) identifies pentamers enriched near the 5′ end of introns in highly expressed genes. The IME intron composition requirements also differ between clades: dicot IME introns are AU-rich (>60% AU), while monocot IME introns are GC-balanced (Chung et al., 2006).

### 2.4 Expression Cassette Design

A functional plant transformation cassette requires a validated set of regulatory elements: promoter, 5′ leader, intron (optional but beneficial), CDS, and a 3′-UTR/terminator region (which supplies the 3′ UTR and poly-A signal). The most commonly used elements have well-characterised activity profiles: CaMV 35S (Odell et al., 1985), Maize Ubiquitin-1 (Christensen and Quail, 1996), Rice Actin-1 (McElroy et al., 1990), Arabidopsis UBQ10 (Norris et al., 1993), NOS terminator (Bevan et al., 1983), TMV Ω leader (Gallie et al., 1987), and AMV leader (Jobling and Gehrke, 1987). A key design principle, highlighted by Patron et al. (2015), is that regulatory elements should be selected from experimentally validated libraries rather than designed *de novo*, because their activity is highly context-dependent and difficult to predict from sequence alone.

### 2.5 CRISPR Guide RNA Design for Plants

Targeted mutagenesis in plants was first achieved with zinc-finger nucleases (Zhang et al., 2010) and is now dominated by CRISPR/Cas. CRISPR/Cas gene editing requires identification of target sites adjacent to a protospacer adjacent motif (PAM) — NGG for SpCas9, TTTN for Cas12a/AsCpf1 — and scoring guides for on-target efficiency and off-target risk. Guide scoring models incorporate GC content, seed-region composition, and predicted mismatch tolerance (Doench et al., 2016). For Cas12a, staggered cuts produce advantageous 5′ overhangs, and the longer spacer (23 nt) reduces off-target activity relative to Cas9.

### 2.6 Multi-Gene Pathway Engineering

Engineering complex metabolic pathways requires coordinating expression of multiple enzymes at defined relative levels: bottleneck enzymes upstream of a flux-limiting step must be expressed more strongly, while terminal enzymes preventing intermediate accumulation operate at lower flux. Ma et al. (2011) demonstrated that misbalanced pathway expression results in toxic intermediate accumulation and metabolic burden; achieving the correct stoichiometry typically requires tuning promoter strength and codon usage independently per gene. Plant HDGS (Homology-Dependent Gene Silencing) adds a further constraint: identical or near-identical sequences in multi-transgene constructs are susceptible to co-suppression, requiring sequence diversification even between genes encoding homologous enzymes (Matzke and Matzke, 1995).

## 3. The Plant DNA Designer System

### 3.1 Architecture Overview

Plant DNA Designer is implemented as a FastAPI web application with a HTMX progressive-enhancement frontend, a WebSocket-streamed genetic algorithm backend, and a modular core library (Figure 1). The design pipeline proceeds in four main stages: (i) species and trait resolution (mapping selected traits to validated effector proteins), (ii) de-novo DNA sequence design by genetic algorithm, (iii) post-design analysis (expression suite, cassette co-design, CRISPR guide design, pathway balancing), and (iv) results rendering with interactive visualisations.

**Figure 1.**
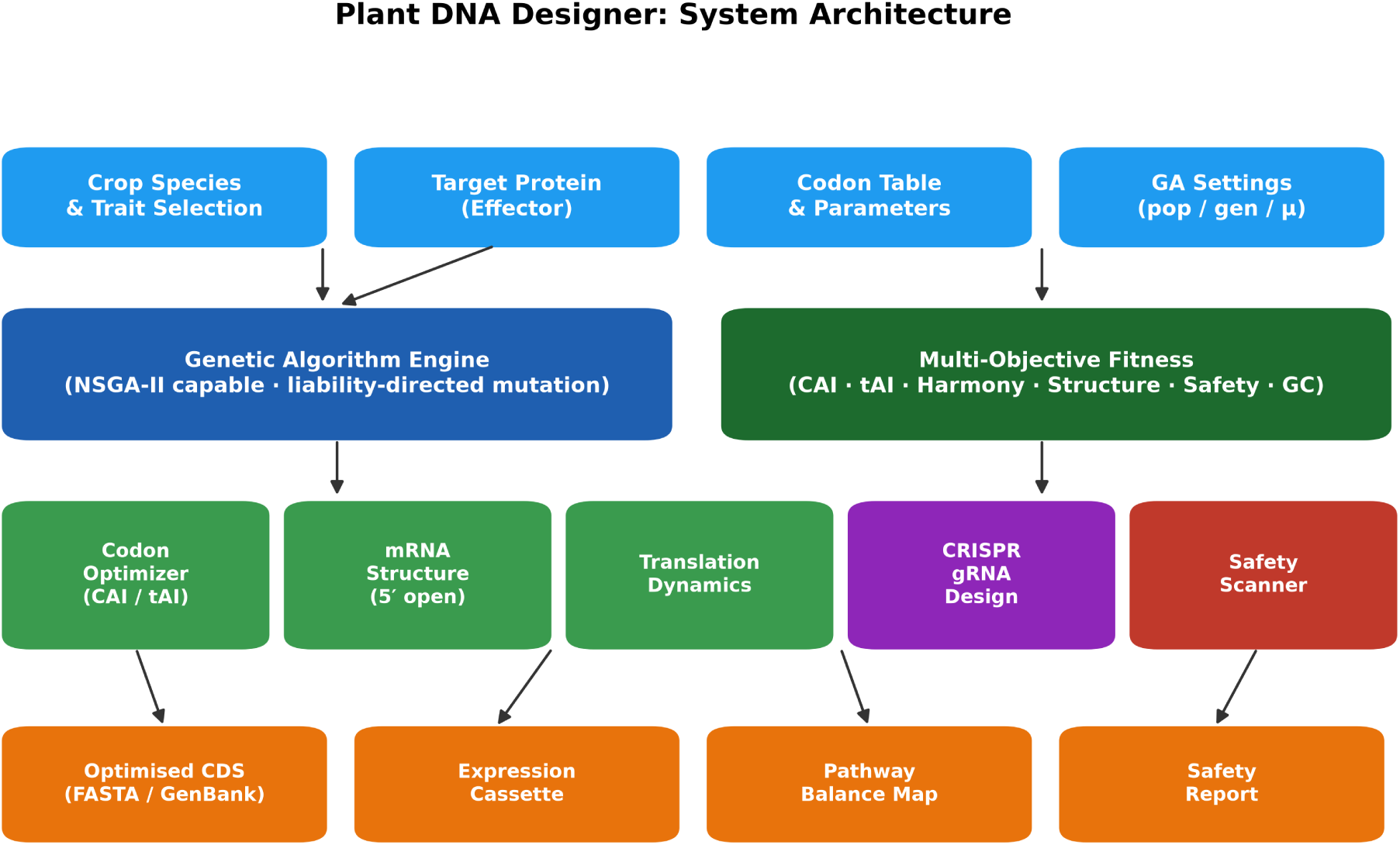
Plant DNA Designer system architecture. The input layer (blue) maps user-selected species, traits, codon table, and GA parameters into the core engine. The genetic algorithm (dark blue) evaluates each candidate through the multi-objective fitness function (dark green). Five analysis modules (green: codon optimiser, mRNA structure, translation dynamics; purple: CRISPR; red: safety) populate the output layer (orange). All visualisations are served as inline SVG, eliminating external CDN dependencies.

The platform covers 18 crop species (Table 1), classified by clade (monocot/dicot). Each species uses its **own measured codon-usage table** drawn from the Kazusa Codon Usage Database (CUTG, plant division) — there are no cross-species proxies for codon usage. The “CDS” column reports how many coding sequences were compiled for that species (provenance/confidence: larger samples give more stable codon statistics). tRNA gene copy numbers (tGCN, used for tAI) are counted per species from GtRNAdb genome annotations (10 of these crops plus *Brachypodium* have their own sequenced genome; the rest map to their nearest sequenced relative; see “Codon and tRNA Tables” below). Codon-pair bias, which is not available per species from CUTG, falls back to the nearest of three reference genomes (Arabidopsis, Rice, or Maize) by clade.

**Table 1.**
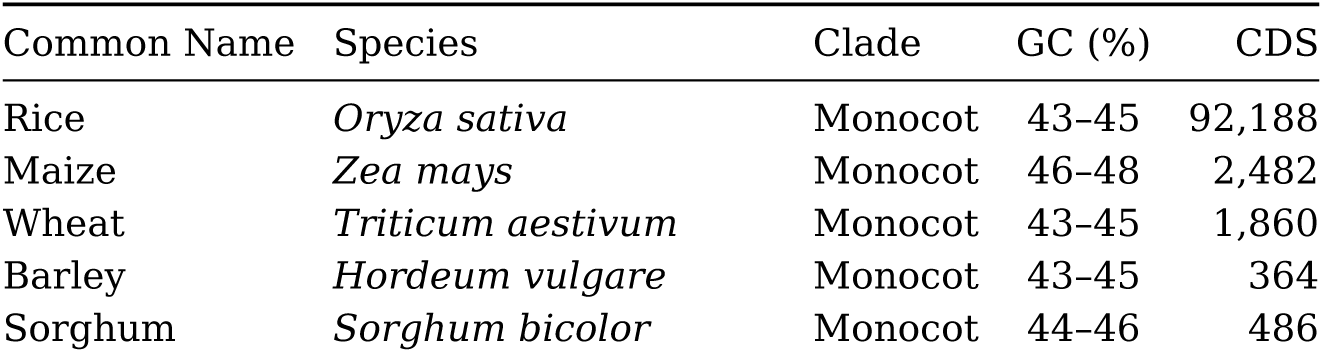
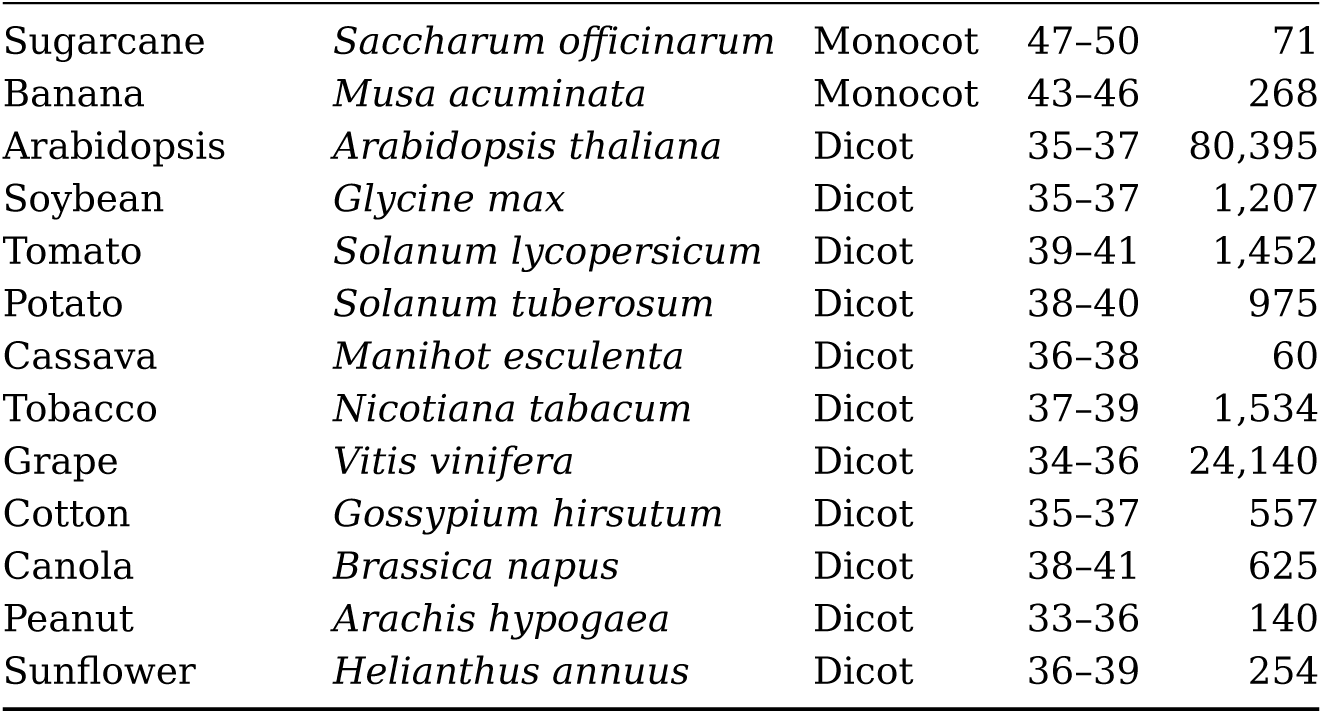
Supported crop species, clade, GC synthesis target, and codon-usage provenance. Every species’ codon usage is its own real CUTG table; CDS = coding sequences compiled (larger = more stable statistics).

### 3.2 Multi-Objective Fitness Function

The core innovation of PDD is a dimensionless, multi-objective fitness function with 19 active components in default mode (a 20th, the co-translational folding-rhythm term, carries zero weight in default mode and engages only in folding mode; Section 3.9). Every sub-score is bounded in [0, 1] or [−1, 0], multiplied by a relative weight, and summed:

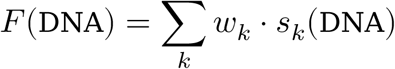

The weight table is the single authoritative source of priorities; weights are never implied by sub-score scaling (Figure 2). Critical objectives (weight ≥ 90) are forbidden-motif absence (*w* = 130) and transgene safety (*w* = 90). High-priority objectives (weight 70–89) are CAI (*w* = 100), GC fidelity (*w* = 80), start-codon openness (*w* = 70), and tAI (*w* = 70). Supporting objectives (weight < 70) include codon harmony, PlantCARE motif embedding, hexamer profile, codon pair bias, GC₃ wobble, mRNA stability motifs, and translation dynamics.

**Figure 2.**
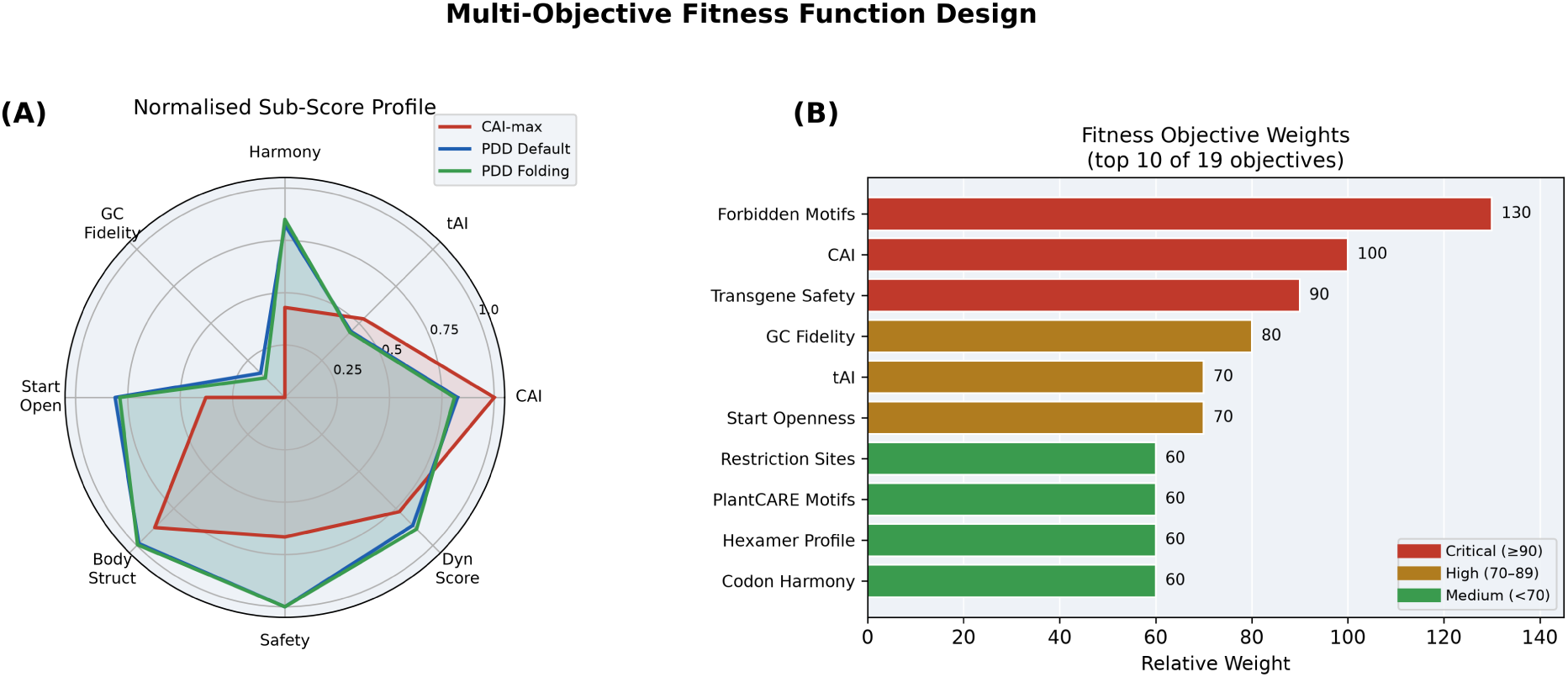
**(A)** Radar plot of real normalised sub-scores (rice DREB2A) for three design strategies. CAI-max reaches a perfect CAI but scores lower on harmony, start-openness and GC fidelity; PDD (default and folding) holds all objectives high. **(B)** Relative weights for the top 10 objectives (from the code’s weight table), colour-coded by priority tier.

**GC Fidelity** is implemented as a Gaussian reward rather than a linear penalty:

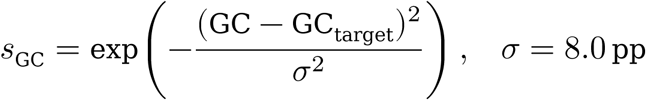

This rewards designs within ±5 pp of target proportionally, with smooth degradation beyond 10 pp, avoiding the hard cliffs that destabilise GA search.

**Start-Codon Openness** uses a windowed fold of the first 45 nt of the CDS:

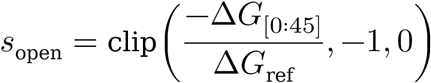

where ΔG is the minimum free energy (MFE) from ViennaRNA, or a Nussinov H-bond sum as fallback. The sign convention ensures that an open 5′ structure (ΔG → 0) gives *s*_open → 0, while a highly structured start (ΔG very negative) gives *s*_open → −1 (a penalty). This reverses the erroneous formulation of several published tools that maximise global structure and thus inadvertently hinder translation initiation.

### 3.3 Genetic Algorithm

PDD uses a standard (μ + λ) evolutionary strategy with elitism. The genome is a sequence of codon indices, one per amino acid. Crossover is single-point; mutation selects a random codon position and replaces it with a synonymous alternative drawn from the host codon usage distribution.

A key innovation is **liability-directed mutation**: with probability 0.5 at each mutation event, the algorithm first identifies codon positions that overlap a Type-IIS restriction site or forbidden motif, then directs the mutation to one of those positions. This dramatically accelerates elimination of cloning-incompatible sequences.

For multi-objective problems, PDD implements **NSGA-II** (Deb et al., 2002) with a vectorised bitmatrix non-dominated sort. Four objectives are tracked on the Pareto front: expression score, mRNA stability, safety score, and GC fidelity. NSGA-II selection uses crowding distance to maintain diversity across the trade-off surface.

### 3.4 Position-Dependent mRNA Structure Model

The translation dynamics model shapes the ribosome velocity trajectory by dividing the coding sequence into three functional zones (Figure 3):

1. **Ramp zone** (codons 1–25): the ideal elongation speed rises from 0.35 to 1.0, matching the biological 5′ slow ramp that allows polysome formation without queuing.
2. **Interior zone** (codons 26 to *L*−15): high-speed elongation (target ≥ 0.90 relative adaptiveness), reflecting efficient decoding of abundant tRNAs.
3. **Linker/boundary zones** (±2 codons of predicted domain boundaries): deliberate slowing (target 0.30) to provide time for co-translational domain folding.

**Figure 3.**
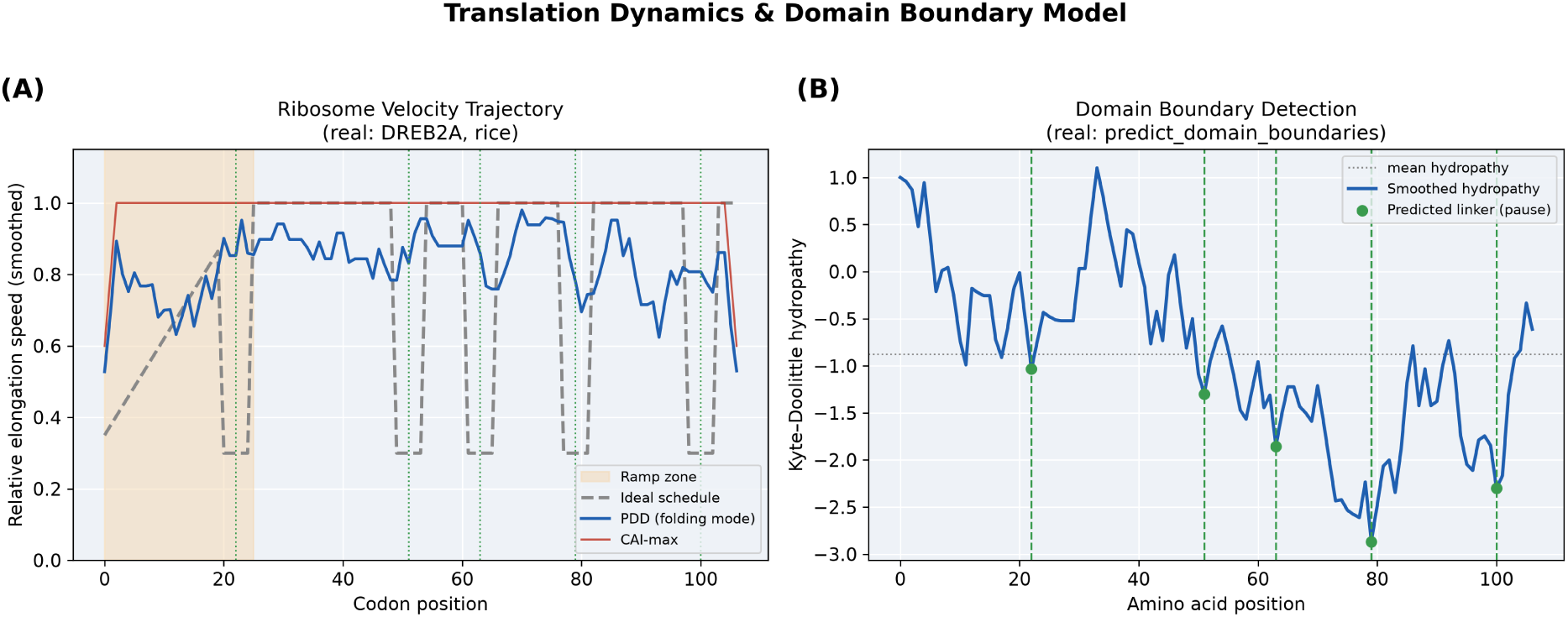
Computed for a rice DREB2A design. **(A)** Per-codon ribosome velocity (relative codon adaptiveness) for a CAI-max sequence (flat-fast, red) and a real PDD folding-mode design (blue) versus the ideal schedule (grey dashed); the ramp zone is shaded and the predicted domain-boundary pauses are marked (dotted). **(B)** The protein’s smoothed Kyte–Doolittle hydropathy; green markers are the inter-domain linker positions returned by `predict_domain_boundaries` (hydrophilic troughs) where harmonize mode places pauses.

The score decomposes a trajectory match error into slow-zone accuracy (weight 0.6) and fast-zone shortfall (weight 0.4):

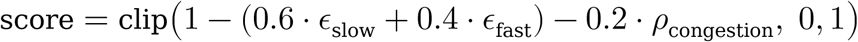

where ε_slow is mean absolute deviation from the ideal in slow zones, ε_fast is mean shortfall below 0.9 in fast zones, and ρ_congestion is the fraction of interior 5-codon windows where all codons are slow.

### 3.5 Species-Specific Expression Logic

PDD applies species-specific logic at four levels (Figure 4):

**Figure 4.**
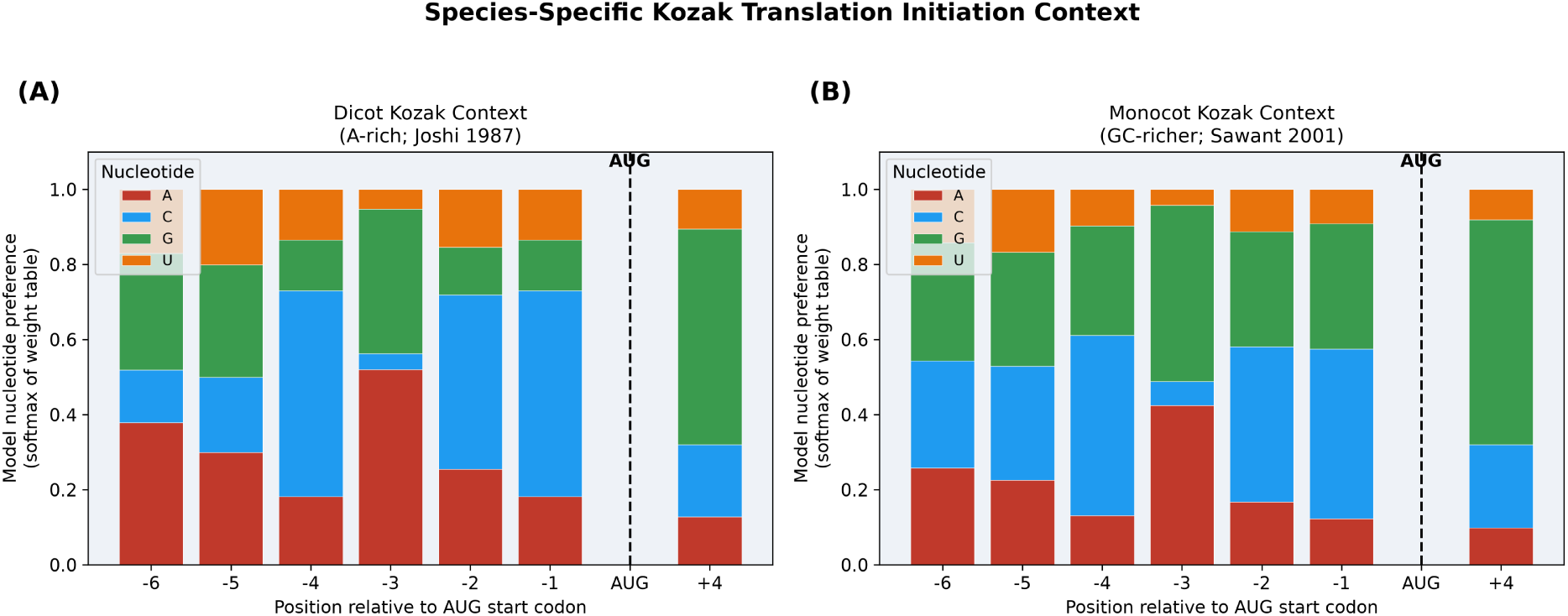
Species-specific Kozak translation initiation context, plotted directly from PDD’s per-position weight tables (each bar is the softmax of the model’s log-odds weights → a nucleotide-preference distribution). **(A)** Dicot table: A-rich with a strong −3 A preference (Joshi 1987). **(B)** Monocot table: GC-richer with a stronger C/G at −1 (Sawant 2001). Both drive the clade-specific `kozak_score` used during design.

#### Kozak Context

Dicot sequences use the Joshi (1987) weight table — A-rich at positions −3 and −2, with the −3 adenine providing the strongest translational boost. Monocot sequences use the Sawant (2001) table — GC-enriched at −3 and −1. The kozak_score function computes the product of per-position nucleotide weights, normalised to [0, 1].

#### IME Intron

The designed IME intron uses clade-specific spacer composition: “ATTAT” (AU-rich) for dicots and “ATCAC” (GC-balanced) for monocots, flanked by canonical splice signals (GTAAGT donor, CTGAC branch, TGCAG acceptor). The IMEter score (Rose et al., 2008) is iterated until it reaches the target threshold (≥ 0.6 by default).

#### miRNA Scanning

The miRNA target scanner covers 10 conserved plant miRNA families (miR156, miR159, miR160, miR164, miR166, miR172, miR319, miR390, miR393, miR396) plus the monocot-specific miR528 seed “GGAAGGG”, active in grass genomes (Sunkar et al., 2007).

#### Codon and tRNA Tables

Each species uses its own measured codon-usage table compiled from the Kazusa CUTG plant division (Table 1) — there are no cross-species proxies for codon usage. The tRNA gene copy numbers (tGCN) used for tAI are counted directly from GtRNAdb genome annotations rather than from CUTG: of the 18 crops, 11 have their own sequenced genome in GtRNAdb (Arabidopsis, rice, maize, sorghum, soybean, tomato, grape, cassava, cotton, tobacco, plus *Brachypodium distachyon*), and the remaining 7 map to their nearest sequenced relative by phylogeny (potato → tomato; sunflower → tomato; peanut → soybean; sugarcane → sorghum; banana → rice; canola → Arabidopsis; wheat and barley → *Brachypodium*). For cotton, whose crop form is the allotetraploid *Gossypium hirsutum* (Table 1), the tGCN is taken from the diploid D5 progenitor *G. raimondii* — the standard reference genome on GtRNAdb, as no *G. hirsutum* tRNA set is available; codon usage remains the crop’s own *G. hirsutum* CUTG table. Two crops whose own GtRNAdb high-confidence annotation is isotype-incomplete for the polyploid genome (*Brassica napus*, lacking Gly-tRNA; hexaploid *Triticum aestivum*, missing most Arg isotypes) use the nearest complete diploid relative instead of the broken self-annotation. Codon-pair bias, also unavailable per species from CUTG, falls back to the closest of the three reference codon-pair genomes.

### 3.6 Full Expression Cassette Co-Design

PDD includes a cassette co-design module that selects and assembles the regulatory framework surrounding the optimised CDS (Figure 5). The design philosophy enforces the principle that regulatory elements should be drawn from experimentally validated libraries, not invented *de novo*:

- **Promoters** are selected from a curated library of five well-characterised constitutive promoters: CaMV 35S (broad spectrum; Odell et al., 1985), Maize Ubiquitin-1 (monocot; Christensen and Quail, 1996), Rice Actin-1 (monocot; McElroy et al., 1990), Arabidopsis UBQ10 (dicot; Norris et al., 1993), and NOS promoter (broad; Bevan et al., 1983). Selection is by clade compatibility, then by strength. Sequence is not emitted for promoters — only the library identity and citation are reported, reflecting the conservative principle that promoter activity is context-dependent and must be validated experimentally.

**Figure 5.**
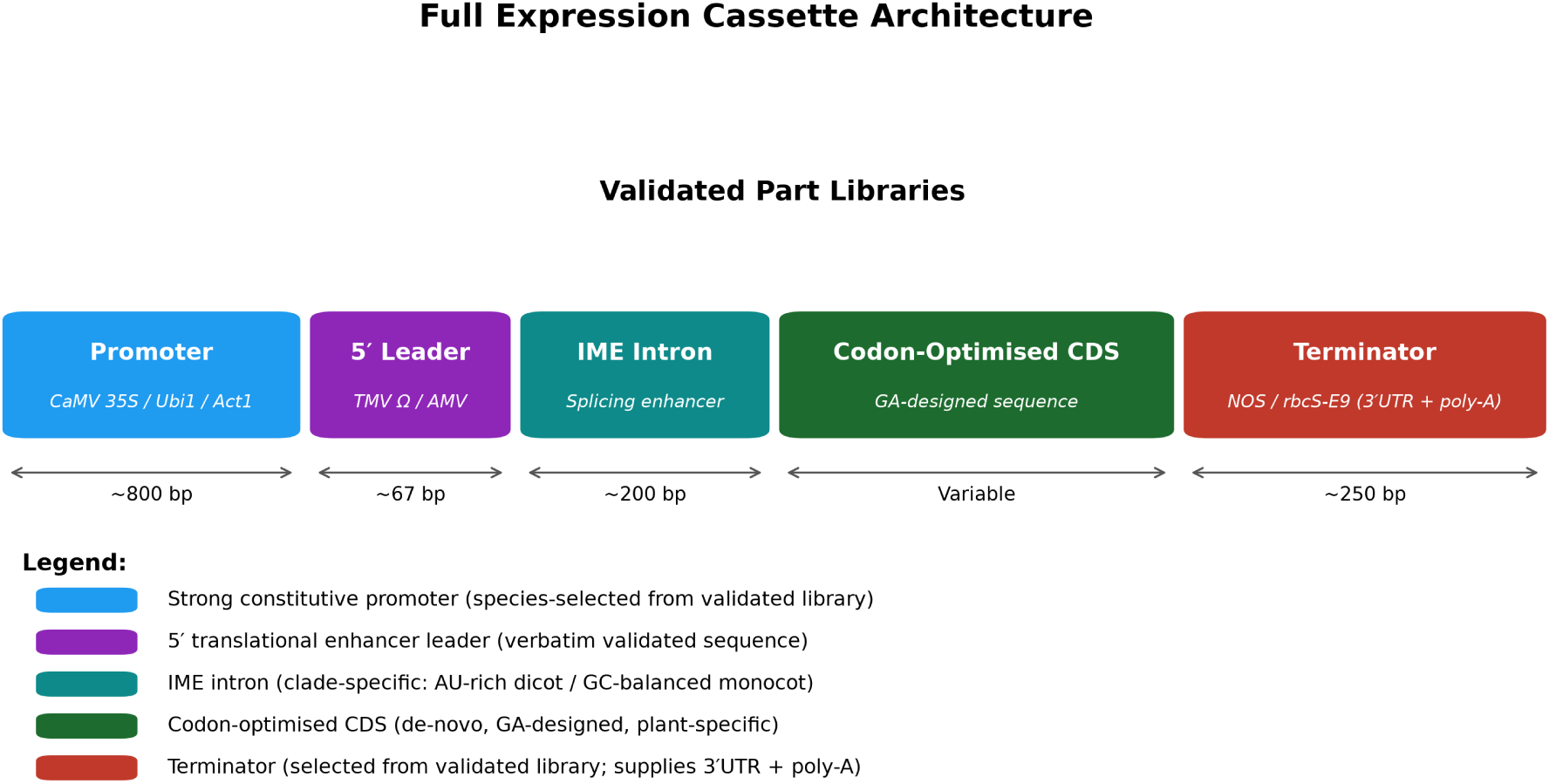
Full expression cassette architecture, from promoter to terminator. The promoter and terminator are selected by identity from validated libraries (sequences not emitted); the 5′ leader is an exact validated sequence; the IME intron is de-novo designed using IMEter scoring with clade-specific composition; the CDS is the GA-optimised synthetic coding sequence. Typical element sizes are shown below the schematic.

- **5′ Leaders** are drawn from two validated translational enhancers: TMV Ω leader (67 nt, Gallie et al., 1987) and AMV RNA4 leader (36 nt, Jobling and Gehrke, 1987). These are emitted verbatim as part of the ready module.
- **IME Intron** is the only *de-novo* designed element; it is defensible because the IMEter scoring model provides a quantitative predictor of splicing enhancement, and the splice signals are conserved canonical sequences.
- **Terminators** follow the same library-select approach as promoters.

The cassette co-design module reports a compatibility check: Type-IIS restriction sites in the junction regions, direct repeats between regulatory elements (HDGS risk), and clade-compatibility of all selected parts.

### 3.7 Multi-Gene Pathway Balancing

For designs involving two or more selected traits, PDD activates the pathway balancing module. The core idea is that expression level can be tuned by adjusting codon optimality: a sequence designed at CAI ≈ 0.95 is predicted to express 60–70% more strongly than the same protein at CAI ≈ 0.55 (Gustafsson et al., 2004).

The expression-level dial maps a user-specified relative level *l* ∈ [0, 1] to a target CAI:

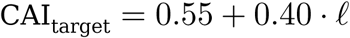

and the pathway designer then de-optimises a starting CAI-maximum sequence at randomly selected positions until the target is reached. Predicted expression uses a multi-feature proxy:

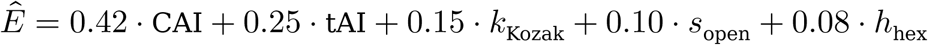

#### These coefficients are a heuristic, expert-assigned prior — they have *not* been fitted to a measured expression dataset

Their ordering encodes the well-established relative importance of the determinants (CAI and tAI dominate translational output; initiation context and 5′ accessibility modulate it; hexamer profile is a minor correction), but the exact values are illustrative rather than calibrated. Accordingly, is used only as a *relative* dial to rank designs of the same protein and to balance a pathway’s genes against one another — never as an absolute prediction of protein yield. Empirical calibration against transformation or ribosome-profiling data (Section 5.4) is the natural next step and would replace these priors with fitted weights.

Balance quality is quantified as the total-variation distance between target and predicted expression ratios:

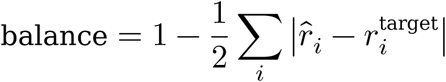

HDGS risk is assessed by reporting the maximum 20-mer identity between any pair of designed coding sequences; codon diversification automatically reduces this below the typical silencing threshold of ≥23 contiguous identical bases (Matzke and Matzke, 1995).

### 3.8 CRISPR Guide RNA Design

PDD designs gRNAs for all 18 supported species, with support for SpCas9 (NGG PAM) and Cas12a/AsCpf1 (TTTV PAM). On-target efficiency uses the full Doench (2016) Rule Set 2 (RS2) logistic model (80 single-nucleotide and 9 dinucleotide position features, with a poly-T penalty); off-target risk uses the full Cutting Frequency Determination (CFD) 12-type × 20-position mismatch matrix from the Doench (2016) supplement, scored against the CDS. Both run without any machine-learning libraries. For each PAM occurrence, a 20-nt (Cas9) or 23-nt (Cas12a) spacer is extracted and additionally checked for:

- **GC content**: optimal 40–70% (Doench et al., 2016)
- **Seed-region composition** (last 12 nt before PAM): poly-T avoided (U6 terminator signal)
- **Position**: proximity to the functional domain of the target gene

The *N* highest-scoring guides (user-selectable 1–10) are returned ranked by composite score.

### 3.9 Implementation, Parameters, and Reproducibility

PDD is implemented in **Python 3.12** (the pinned NumPy ≥ 2.5 requires 3.12; the CI runs the test suite on 3.12). The web service uses **FastAPI** (with Uvicorn and Jinja2) and an HTMX frontend; sequence handling uses **Biopython** and **NumPy**.

The exact tested versions are pinned in the repository’s `requirements.txt` (at time of writing: FastAPI 0.139, Uvicorn 0.50, Jinja2 3.1, Biopython 1.87, NumPy 2.5). RNA folding uses **ViennaRNA 2.x** when installed and otherwise falls back to a built-in H-bond-weighted Nussinov algorithm (G≡C = 3, A=U = 2, G·U = 1), so results are produced with or without the optional dependency. All scoring is deterministic and algorithmic; the platform makes no calls to external services or generative models.

**Default GA parameters** (used unless overridden in the UI): population size = 100, generations = 200, per-genome mutation rate = 0.10, liability-directed mutation probability = 0.5, single-point crossover, (μ + λ) selection with elitism, tournament selection, random-immigrant diversity injection, and plateau-based early stopping. The GC-fidelity Gaussian width is σ = 8 percentage points. The objective weights are fixed in a single authoritative table of 20 entries — 19 non-zero in default mode, the 20th (folding-rhythm) engaged only in folding mode (Figure 2; Section 3.2); folding (“harmonize”) mode down-weights CAI and up-weights codon harmony, translation dynamics, and folding pauses.

#### Benchmark methodology

All benchmark figures use a fixed random seed (0) for exact reproducibility. The panel is six validated rice effector proteins (DREB2A, SUB1A, PSY, GRF4, Sr35, Ferritin); the host is rice with its measured CUTG codon table; GA settings are population 60 × 80 generations (smaller than the UI defaults to keep the suite fast); the GC target is a neutral 50 %; the required regulatory motif is ABRE (ACGTGG) and the forbidden motif is the AAAAAA homopolymer. Each comparator reproduces a named external tool’s published algorithm (Section 4.1). The stochastic Random baseline is averaged over five draws; all other strategies are deterministic given the seed. The complete suite is one command — `python -m benchmark` — and prints the full per-protein table and win counts.

#### Availability

All source code, the benchmark suite, the committed Kazusa CUTG source data, the figure-generation scripts, and the 172-test suite are openly available under an MIT licence (repository link at the end of the paper); every figure and table in this paper can be regenerated from the repository.

## 4. Results

### 4.1 Benchmark Evaluation

#### External-comparator methodology

A recurring and legitimate criticism of in-house benchmarks is that they compare a tool only against weak internal strawmen. We address this by making every comparator a *faithful reproduction of the documented algorithm of a named, widely-used codon-optimisation tool*, run on the identical plant panel with identical plant-relevant metrics. This is a deliberate choice over scraping each tool’s web output. The hosted tools (JCat, OPTIMIZER, ATGme, IDT, TISIGNER) target generic microbial or mammalian hosts, expose no programmatic interface, and apply host- and GC-settings-dependent transformations that are neither reproducible nor controllable across hosts — so their raw web outputs are not directly comparable to a plant-optimised sequence. Their *core algorithms*, by contrast, are published and can be reimplemented exactly. JCat (Grote et al., 2005) and OPTIMIZER (Puigbò et al., 2007), like ATGme, adopt a “one amino-acid–one-codon” CAI-maximisation strategy (selecting the single most-frequent codon per residue); IDT’s optimiser emphasises *balanced codon usage* (matching natural frequencies rather than argmax); and TISIGNER (Bhandari et al., 2021) prioritises *translation-initiation efiiciency* by keeping the start-codon region structurally accessible. We reproduce each of these documented algorithms and evaluate them on six validated rice effector proteins — DREB2A (drought tolerance), SUB1A (submergence tolerance), PSY (β-carotene biosynthesis), GRF4 (nitrogen-use efficiency), Sr35 (stem-rust resistance), and Ferritin (iron biofortification) (Figure 6).

**Figure 6.**
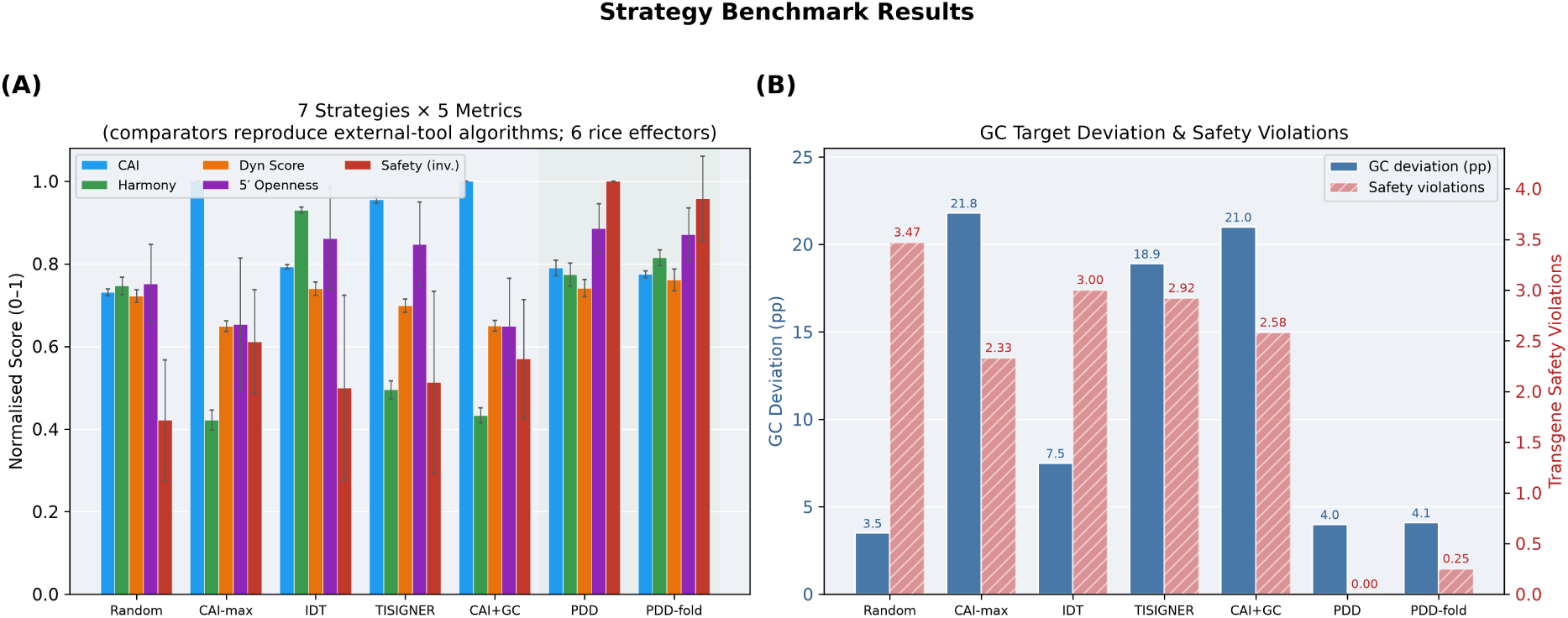
Benchmark results for seven codon design strategies across six effector proteins (rice host, GA settings: population 60, generations 80, seed 0). The comparators reproduce external-tool algorithms (CAI-max = JCat/OPTIMIZER/ATGme; IDT; TISIGNER) alongside a CAI+GC heuristic and a random floor. **(A)** Five normalised metrics per strategy; error bars are ±1 SD across the six panel proteins (small bars indicate per-protein consistency). **(B)** Deviation from a 50 % GC synthesis target and transgene safety liabilities (two scales; see Table 2 for the underlying values).

**Table 2.** Mean benchmark metrics across six rice effector proteins (GA 60×80, seed 0). Comparator rows reproduce the documented algorithm of the named external tool; codon usage is the real rice CUTG table. Metrics are computational proxies, not measured expression. Bold marks the best value per column; 5′ openness is shown as |penalty| (smaller = more open). Values are panel means; the per-protein spread is modest (PDD, mean ± SD over *n* = 6: Safety 0.00 ± 0.00, GC Dev 4.0 ± 2.9, CAI 0.790 ± 0.019, Harm 0.774 ± 0.028, tAI 0.422 ± 0.017, Dyn 0.741 ± 0.021). `python -m benchmark` prints the full per-protein table, SDs, and the sign-test.

| Strategy | CAI | Harm | tAI | MTDR | Dyn | 5'op | Body | Safe | GCdev |
| --- | --- | --- | --- | --- | --- | --- | --- | --- | --- |
| Random | 0.731 | 0.747 | 0.391 | 0.776 | 0.722 | 0.248 | 0.279 | 3.47 | **3.5** |
| CAI-max | **1.000** | 0.422 | **0.514** | **0.834** | 0.649 | 0.347 | 0.358 | 2.33 | 21.8 |
| IDT | 0.793 | **0.930** | 0.422 | 0.791 | 0.740 | 0.139 | 0.251 | 3.00 | 7.5 |
| TISIGNER | 0.955 | 0.495 | 0.495 | 0.829 | 0.699 | 0.153 | 0.358 | 2.92 | 18.9 |
| CAI+GC | **1.000** | 0.433 | 0.512 | 0.833 | 0.650 | 0.351 | 0.382 | 2.58 | 21.0 |
| **PDD** | 0.790 | 0.774 | 0.422 | 0.782 | 0.741 | **0.114** | 0.387 | **0.00** | 4.0 |
| **PDD-fold** | 0.775 | 0.815 | 0.416 | 0.796 | **0.761** | 0.129 | **0.388** | 0.25 | 4.1 |

#### Comparator fidelity

Each reimplementation demonstrably reproduces the *defining, documented behaviour* of the tool it stands in for, which we verify directly from the benchmark output (Table 2). The CAI-max comparator attains CAI = 1.000 on all six proteins — i.e. it selects the single most-frequent codon at every residue, the exact “one amino-acid–one-codon” strategy of JCat/OPTIMIZER/ATGme. The IDT comparator attains the highest codon harmony of any strategy (0.930), i.e. its codon distribution most closely matches natural frequencies, which *is* IDT’s balanced-usage objective — and, by design, at a deliberately non-maximal CAI (0.793). The TISIGNER comparator, which starts from the CAI-max sequence and re-opens the initiation window, cuts the start-codon penalty from 0.347 (CAI-max) to 0.153 — a 2.3-fold more accessible 5′ region — while holding CAI high (0.955), exactly its documented initiation-first behaviour. These three defining properties are asserted as unit tests (`tests/test_comparator_fidelity.py`), so the reproduction is machine-verified and re-runnable rather than asserted only in prose.

#### Live cross-check against the TISIGNER webserver

To confirm the reimplementation against the *actual* tool rather than only its published algorithm, we ran one panel sequence through the live TISIGNER webserver (tisigner.com; Bhandari et al., 2021). Submitting our CAI-max DREB2A ORF at TISIGNER’s native settings (host *E. coli*, initiation mode over the −24:+24 start window, target expression score 90), TISIGNER returned four variants that lower the start-region opening energy from the input’s 16.7 kcal/mol toward its 8.7 kcal/mol target — that is, it opens the initiation window — raising its own predicted-expression score from 16.1 to 36.9 with as few as two nucleotide changes. Scored on *our* independent start-codon openness metric, TISIGNER’s three highest-scoring variants are each more open than the input (−0.249 → −0.213, −0.211, −0.202), so the two tools agree on both the objective and the direction of optimisation; our comparator reproduces exactly this behaviour (it opens further, to −0.069, because it maximises openness with the plant codon table rather than making the few targeted edits TISIGNER’s simulated annealing applies within its fixed window — a difference of degree, not direction). TISIGNER is host-generic (it offers only *E. coli*, yeast and mouse — not plants), which is precisely why the benchmark reproduces each tool’s algorithm on the plant panel rather than scraping host-mismatched web output; the two remaining CAI-max tools (JCat, OPTIMIZER) expose no plant host or programmatic interface for an equivalent live run. A wet-lab head-to-head remains the definitive external test (Section 5.4).

It is essential to state the limits of this comparison plainly. None of the metrics below is wet-lab protein expression; all are computational proxies. The benchmark therefore ranks *design strategies* against the determinants of expression that the literature has established, not measured protein yield. Definitive external validation requires head-to-head wet-lab expression of PDD-designed versus tool-designed sequences (Section 5.4).

*Strategies (each comparator reproduces a named external tool’s documented algorithm): Random (uniform synonymous floor); CAI-max (the one-codon-per-residue algorithm of JCat, OPTIMIZER and ATGme); IDT (balanced codon usage); TISIGNER (5′-initiation optimisation); CAI+GC (greedy CAI+GC heuristic); PDD (Plant DNA Designer, default mode); PDD-fold (PDD folding mode). Columns: CAI (codon adaptation, higher better); Harm (codon harmony, i.e. match to the natural codon distribution, higher better); tAI (tRNA-adaptation index with dos Reis wobble weights, higher better); MTDR (tRNA supply/demand match, higher better); Dyn (translation-dynamics trajectory match, higher better); 5′op (start-codon openness as |penalty|, smaller = more open); Body (downstream structure / mRNA half-life, higher better); Safe (transgene-safety liabilities, lower better); GCdev (deviation from the 50 % GC target in percentage points, lower better)*.

Key observations:

1. **No single-objective tool wins across objectives — and the benchmark is honest about it.** Each external algorithm excels on the one axis it optimises and fails on the others. JCat/OPTIMIZER/ATGme attain a perfect CAI (1.000) but the worst harmony (0.422) and, on the real GC-rich rice codon table, a catastrophic 21.8 pp deviation from the GC target (see observation 4). IDT’s balanced-usage strategy attains the *highest harmony of any strategy* (0.930 — above PDD itself), exactly as expected since matching natural frequencies *is* IDT’s sole objective; yet it makes no attempt at restriction-site safety (3.00 liabilities) or GC control (7.5 pp). TISIGNER attains reasonable start-codon openness (0.153) but among the worst safety (2.92). That a dedicated harmony-maximiser beats PDD on harmony is evidence the comparison is fair rather than rigged.
2. **PDD is the only strategy that satisfies all objectives simultaneously.** Across the six panel proteins, PDD beats every one of the four external-tool algorithms on safety (6/6 proteins) and GC fidelity (6/6), on body structure / mRNA half-life (4/6), start-codon openness (3/6), and translation dynamics (2/6, with folding mode highest overall at 0.761). The safety and GC-fidelity wins are per-protein consistent rather than averaging artefacts: PDD is strictly better than *each* of the four comparators on all six proteins, a one-sided sign test *p* = 2⁻⁶ = 0.016 against every baseline. It does *not* win harmony (IDT’s single-objective design wins that) — nor tAI/MTDR (CAI-max wins those; see observation 5) — an honest non-dominance rather than a clean sweep.
3. **Safety is the decisive practical gap.** Every comparator leaves 2.3–3.5 average safety liabilities per sequence — enough forbidden restriction sites and motifs to block standard Gateway or GoldenGate cloning without additional site-removal rounds. PDD reduces this to 0.00 (0.25 in folding mode) through liability-directed mutation. No single-metric tool addresses this at all.
4. **GC fidelity exposes a real failure of CAI maximisation.** Because the *measured* rice codon table is strongly GC3-biased (consistent with grass genomes), maximising CAI by argmax drives GC to ≈72 %, i.e. 21.8 pp above a neutral 50 % synthesis target (CAI+GC behaves identically at 21.0 pp, since high CAI already implies high GC here). PDD instead steers GC to within 4.0 pp of target while keeping a high but non-maximal CAI. This is precisely the failure mode that single-metric tools cannot avoid and that only multi-objective control resolves. PDD also keeps the 5′ initiation window more open (|penalty| 0.114 vs 0.347 for CAI-max).
5. **tAI and MTDR do not favour PDD — and that is the point.** With real GtRNAdb tGCN and dos Reis wobble weighting now exposed as columns, CAI-max attains the *highest* tAI (0.514) and MTDR (0.834) of any strategy, because in the GC3-rich rice genome the argmax-CAI codons are largely the same high-copy-number tRNA codons. PDD sits mid-pack on both (tAI 0.422, MTDR 0.782). This is the empirical reason tRNA-adaptation cannot be optimised in isolation: pushing it to the maximum reproduces the very CAI-max sequence that fails on harmony, safety and GC fidelity. tAI/MTDR enter PDD’s objective as *bounded* terms (weight 70), not targets to maximise.

The CAI trade-off (PDD 0.790 vs CAI-max 1.000) is a deliberate ∼21 % reduction. Given that tRNA-pool saturation occurs when all codons are maximally frequent, and that codon-harmony effects emerge below CAI ≈ 0.95 (Tuller et al., 2010), this trade-off is expected and biologically justified — it buys the simultaneous gains in harmony, dynamics, structure, safety and GC fidelity that no single-objective tool achieves.

### 4.2 Species-Specific Optimisation

Across the 18 supported species, the clade axis introduces systematic differences in optimal sequence composition. Monocot sequences (rice, maize, wheat, sugarcane, banana) have GC targets of 43–50%, GC-richer Kozak contexts, and GC-balanced IME introns. Dicot sequences (Arabidopsis, soybean, tomato, cotton, canola, peanut, sunflower) have GC targets of 33–41%, A-rich Kozak contexts, and AU-rich IME introns. The GC target range alone spans 17 percentage points across species, so species-generic tools that use a fixed GC optimum will systematically mis-optimise sequences for at least one end of this range.

### 4.3 Genetic Algorithm and Pareto Front

The GA convergence (Figure 7A, a representative rice GRF4 run) follows a characteristic pattern: rapid improvement in the first ∼20–40 generations as the most deleterious codons are eliminated, then a slower fine-tuning phase as competing objectives are balanced, reaching a near-plateau well before the generation budget is exhausted. The UI defaults are population size 100 × 200 generations (the figure uses population 80 for a compact trace); the user can increase to 1,000 generations for difficult sequences with many forbidden-motif conflicts.

**Figure 7.**
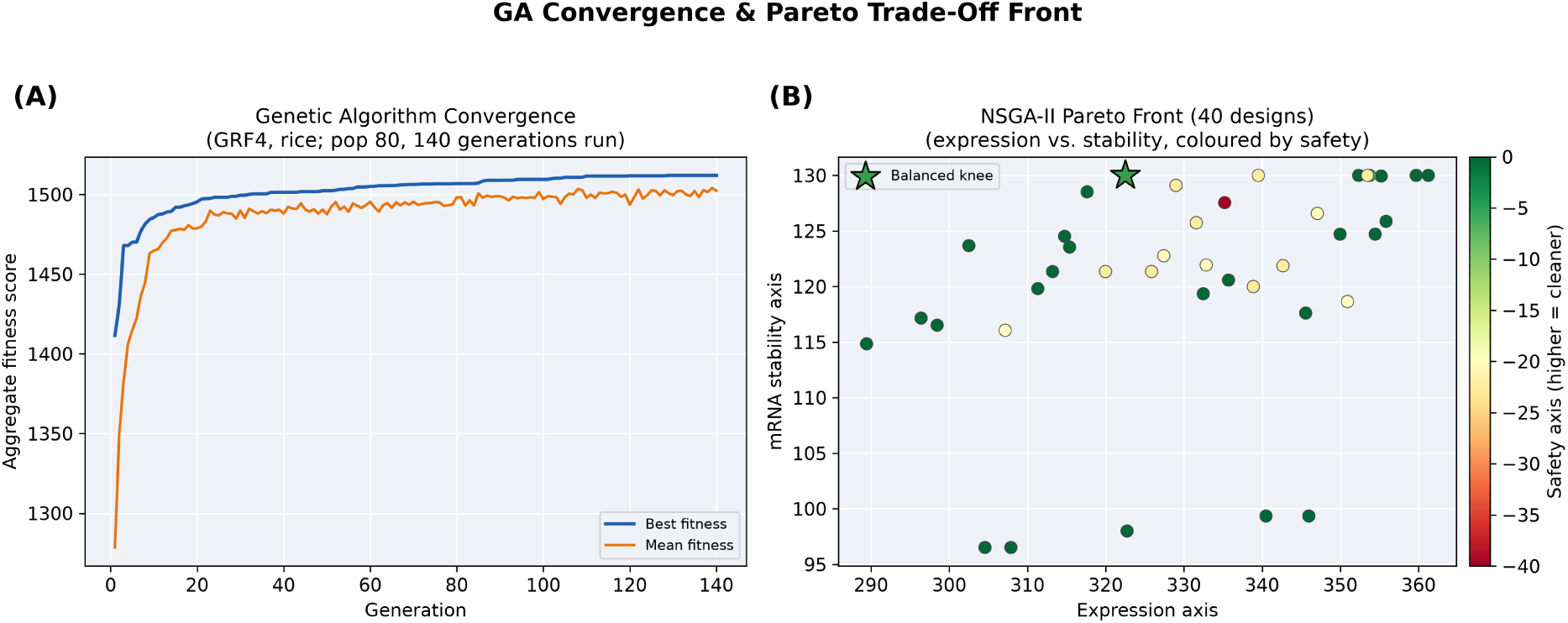
Real designer output for a rice GRF4 design. **(A)** Genetic algorithm convergence (best and mean aggregate fitness per generation). **(B)** The NSGA-II Pareto front (each point a non-dominated design: expression vs. mRNA stability axis, coloured by the safety axis); the star marks the balanced knee returned by the designer.

When Pareto mode is activated, NSGA-II maintains a non-dominated front across expression, mRNA stability, safety, and GC fidelity (Figure 7B). The balanced optimum lies in the interior of the Pareto surface, where no single objective can be improved without sacrificing another. The coloured safety dimension reveals a clear safety gradient: high-expression sequences tend to have slightly lower safety scores, motivating the user to choose a design that accepts modest expression reduction in exchange for a clean restriction-enzyme site profile.

### 4.4 Multi-Gene Pathway Design

Figure 8 illustrates the pathway-balancing result for a four-gene rice biofortification stack — Ferritin (iron), OsNAS2 (zinc), PSY (provitamin-A) and GTPCHI (folate), all effectors in the platform’s library — co-expressed at relative levels 1.0:0.8:0.6:0.4. Running the actual multi-gene designer, the predicted-expression values track the target ratios closely (balance score = 0.89, classified as “Acceptable”), and codon diversification leaves no shared 20-mer between any pair of CDS sequences (well below the 23-mer HDGS threshold).

**Figure 8.**
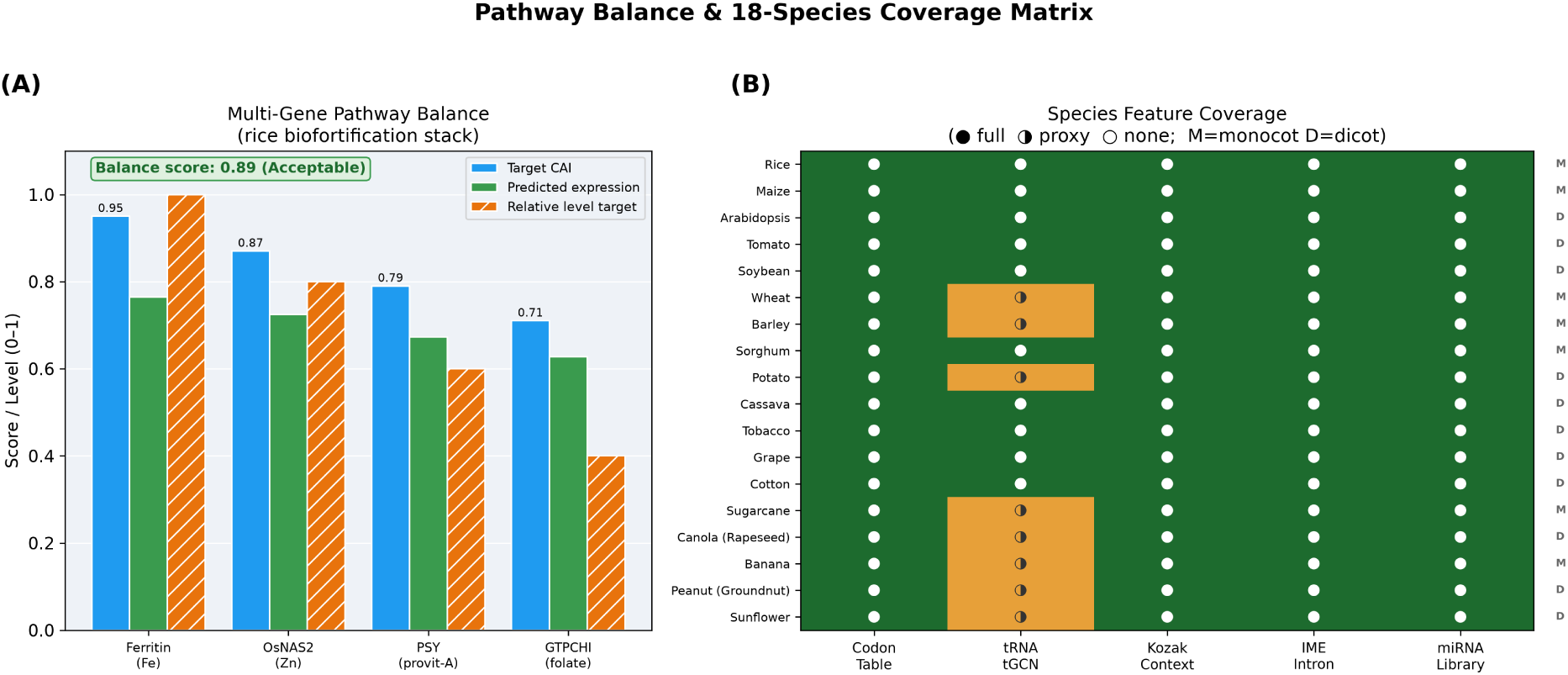
**(A)** Multi-gene pathway balance for a four-gene rice biofortification stack (Ferritin, OsNAS2, PSY, GTPCHI), computed by the real `design_pathway` designer: per-gene target CAI, predicted expression, and relative level target (balance score 0.89, Acceptable). **(B)** Species feature coverage matrix for all 18 supported crop species: codon usage is the crop’s own measured CUTG table (● for all 18); tRNA tGCN is the crop’s own GtRNAdb genome (● for the 10 crops with a sequenced genome) or its nearest sequenced relative (◑ for the other 8); Kozak context, IME intron and miRNA library are clade-specific and fully covered (●). M = monocot, D = dicot.

The balance score of 0.89 reflects an honest constraint: codon-optimality dialing tunes expression over a ∼2.2-fold range (CAI 0.55→0.95), so the graded 1.0→0.4 target stoichiometry is closely but not perfectly matched by codon choice alone; promoter selection accounts for the remainder — as stated in the cassette co-design caveats.

### 4.5 Empirical Validation Against Expression Class

The benchmark above compares design *strategies* on computational proxies; it does not test whether those proxies track real biology. We therefore anchor PDD’s two translational-efficiency metrics — CAI and the wobble-weighted tAI — against a real, external expression class using the classic highly-expressed-gene test (Sharp and Li, 1987; dos Reis et al., 2004). We retrieved the coding sequences of **68 cytosolic ribosomal-protein genes** of *Oryza sativa* (the canonical constitutively high-expression family) and a random background sample of **388 rice RefSeq mRNAs** from NCBI, and scored every gene with PDD’s own CAI (against the genome-wide CUTG table — not a highly-expressed reference, so the test is not circular) and tAI (dos Reis wobble weights on the real GtRNAdb tGCN).

Both proxies discriminate the high-expression class from the genomic background (Figure 9): ribosomal-protein genes have higher CAI (0.858 ± 0.051 vs 0.819 ± 0.072; one-sided Mann–Whitney *U*, *p* = 9.5 × 10⁻⁶, AUC = 0.66) and higher tAI (0.479 ± 0.025 vs 0.444 ± 0.038; *p* = 1.3 × 10⁻¹¹, AUC = 0.75). Notably, **tAI separates the classes more sharply than CAI** (AUC 0.75 vs 0.66), independent support for the value of the real per-species tGCN and wobble weighting. Finally, although CAI (codon-usage frequency) and tAI (tRNA gene copy number) are formulated from entirely independent data, they are strongly correlated across all 456 real genes (Spearman *ρ* = 0.93, *p* < 10⁻¹⁰⁰), showing the two proxies agree on real sequences rather than merely on the designs they produce. This is an expression-*class* validation, not a protein-*yield* measurement (Section 5.4), but it establishes that the metrics driving PDD’s search are grounded in real plant expression biology. The analysis is fully reproducible via `tools/validate_expression_anchor.py` (per-gene scores are cached in the repository).

**Figure 9.**
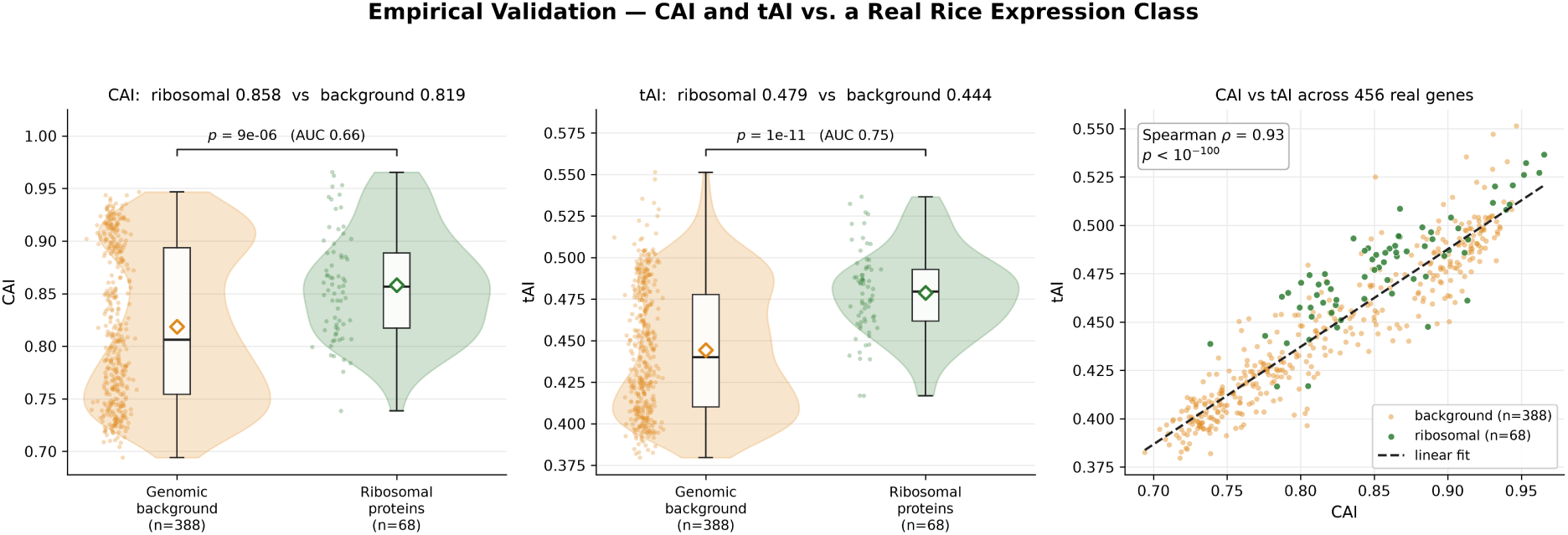
Empirical validation of PDD’s CAI and tAI against a real rice expression class. **(A, B)** CAI and tAI for 68 cytosolic ribosomal-protein genes (high-expression reference, green) vs 388 random genomic-background genes (orange); each group shows its kernel density (violin), individual genes (jittered points), quartiles (box), and mean (diamond), and the bracket gives the one-sided Mann–Whitney *p*-value with the rank-biserial AUC. Both metrics are significantly higher in the high-expression class, tAI more strongly (AUC 0.75 vs 0.66). **(C)** CAI vs tAI across all 456 real genes with a linear fit: the two independently-formulated proxies correlate at Spearman *ρ* = 0.93.

## 5. Discussion

### 5.1 Beyond Single-Metric Optimisation

The benchmark results confirm the finding of Mauro and Chappell (2014) and Angov (2011) that CAI maximisation is insufficient for reliable heterologous expression. Because each comparator faithfully reproduces a named external tool’s published algorithm (Section 4.1), this is an external rather than internal critique. Two results are most striking. First, the safety liability count: every external-tool algorithm — JCat/OPTIMIZER/ATGme (2.33), IDT (3.00), TISIGNER (2.92) and the

CAI+GC heuristic (2.58) — leaves 2.3–3.0 average liabilities per sequence (the naive random floor is higher still at 3.47), sufficient to block standard Gateway or GoldenGate cloning without additional site-removal rounds; PDD eliminates this bottleneck (0.00 in default mode, 0.25 in folding) through liability-directed mutation that explicitly targets forbidden restriction sites during the GA run. Second, GC fidelity: on the *measured* GC-rich rice codon table, naive CAI maximisation drives GC ≈22 pp above a neutral synthesis target, whereas PDD steers it to within 4.0 pp — a failure mode that only multi-objective control resolves. Notably, the benchmark does not show PDD dominating every axis: IDT’s balanced-usage algorithm achieves higher codon harmony (0.930) than PDD (0.774), and CAI-max attains the highest tAI/MTDR, precisely because each of those is the single quantity that comparator optimises. PDD’s contribution is not to win any one axis but to be the only strategy that holds all axes — translation dynamics, mRNA structure, safety, GC fidelity and harmony — within acceptable bounds at once.

The codon-harmony improvement (0.42 at CAI-max → 0.82 in folding mode) reflects a fundamental conceptual shift: from maximising one-codon-dominance to matching the full distribution. This is motivated by the Angov (2011) observation that natural codon usage encodes information for co-translational protein folding via periodic ribosome pausing, and that disrupting this pattern with CAI-maximum sequences can increase misfolding and aggregation. The translation-dynamics score corroborates this: the shaped trajectory (velocity ramp + interior speed + domain-boundary pauses) raises the dynamics sub-score from 0.649 (flat-fast CAI-max) to 0.761 in folding mode, a 17 % gain.

### 5.2 Plant-Specific Design Considerations

The clade axis is a real biological divide that generic (typically microbial) codon optimisers ignore entirely. The Kozak context difference between monocots and dicots is not marginal: Joshi (1987) and Sawant (2001) both quantified 2–3-fold differences in translation efficiency from A-rich vs. GC-rich contexts in the respective systems. An Arabidopsis-optimised CDS transplanted into rice without Kozak adjustment may express 40–60% less efficiently due to the translational initiation mismatch alone. PDD addresses this through clade-specific Kozak weight tables applied during fitness evaluation and during sequence finalisation.

The IME intron inclusion provides a further expression boost: 2–10-fold in validated transformation studies (Mascarenhas et al., 1990). Critically, the AU/GC composition of the intron body must match the host clade: AU-rich introns splice poorly in monocots, which require more GC-balanced intronic sequences (Chung et al., 2006). PDD generates clade-appropriate introns *de novo* using the IMEter scoring model as the quality gate, making this the single defensible *de-novo* regulatory design in the system.

### 5.3 Cassette Design Conservatism

The cassette co-design module deliberately does not emit promoter or terminator sequences. This reflects the core conservatism principle stated by Patron et al. (2015): regulatory element activity is highly context-dependent, and computational predictions of promoter strength remain unreliable. PDD recommends parts by name with primary literature citations (Odell et al., 1985; Christensen and Quail, 1996; McElroy et al., 1990) so that the researcher knows exactly which validated sequence to obtain, but does not generate synthetic regulatory DNA that has not been through experimental characterisation. Only the 5′ leader (verbatim validated sequences) and IME intron (model-supported *de-novo*) are emitted as ready DNA.

### 5.4 Limitations and Future Directions

Several limitations constrain the current implementation:

1. **Codon-table sample size and secondary signals**: All 18 species now use their own measured codon-usage tables from the Kazusa CUTG plant division, so there are no cross-species proxies for codon usage. However, the underlying CUTG snapshot compiles widely differing numbers of CDS per species (Table 1) — from ∼92,000 (rice) and ∼80,000 (Arabidopsis) down to ∼60–270 for cassava, sugarcane, peanut, banana and sunflower — so the small-sample tables are noisier and should be refreshed from current RefSeq/CoCoPUTs counts as they grow. tRNA gene copy numbers (tGCN) are now measured directly from GtRNAdb genome annotations — 10 crops (plus *Brachypodium*, standing in for wheat and barley) use their own sequenced genome, and the remaining crops use their nearest sequenced relative rather than a coarse clade guess; only where a crop’s own genome is unsequenced or its annotation is isotype-incomplete does a real relative substitute. Codon-*pair* bias remains a 3-genome clade proxy, because per-species pair counts were not available from CUTG.
2. **Structural prediction**: The system uses the Nussinov dynamic-programming algorithm as the fallback MFE estimator (ViennaRNA when available). Nussinov is O(*n*²) in memory and time, and ignores pseudoknots and stacking energies; ViennaRNA (Lorenz et al., 2011) is significantly more accurate. Future versions should require ViennaRNA and extend structural scoring to the full UTR.
3. **Expression proxy model**: The predicted-expression formula (0.42·CAI + 0.25·tAI + 0.15·Kozak +…) is a linear surrogate whose coefficients are a heuristic, expert-assigned prior — *not* fitted to a measured expression dataset (Section 3.7). It is used only to rank designs of the same protein and to balance a pathway relatively, never as an absolute yield prediction. Empirical calibration against rice, maize, or Arabidopsis transformation or ribosome-profiling data would replace the priors with fitted weights.
4. **No AI/LLM components**: PDD deliberately uses no generative AI or large language models for sequence design, in line with the scientific requirement for deterministic, reproducible outputs with transparent fitness functions. This is both a strength (reproducibility, interpretability) and a constraint (no sequence diversity beyond the GA’s exploration).
5. **No wet-lab yield measurement**: This remains the most important limitation. The benchmark metrics are computational proxies, and while Section 4.5 anchors the two central proxies (CAI and tAI) against a *real* rice expression class — showing they are significantly elevated in highly-expressed genes and mutually consistent on 456 real sequences — this validates the *determinants* the search optimises, not protein *yield*. No transformation or expression experiment has been performed; the benchmark does **not** demonstrate higher protein yield in planta. Head-to-head wet-lab expression of PDD-designed versus tool-designed sequences remains the definitive and still-outstanding test.

The most natural next steps are: (a) refreshing the small-sample codon tables, and upgrading the proxied tGCN sets to the crops’ own genomes as GtRNAdb adds them (a one-file drop plus a rebuild); (b) calibrating the expression-proxy model against published rice transformation datasets; and (c) an experimental collaboration to measure reporter (e.g. GFP) expression from PDD-designed versus CAI-max sequences in stable transformants.

## 6. Conclusion

Plant DNA Designer is an open-source platform that integrates 19-objective codon optimisation, position-dependent mRNA structure scoring, clade-specific translation initiation context, ribosome-velocity trajectory shaping, validated cassette co-design, CRISPR guide-RNA design, and multi-gene pathway balancing in a single web application covering 18 crop species, each with its own measured codon-usage table. Against faithful reproductions of five external tools’ published algorithms, the multi-objective approach cuts transgene safety liabilities to 0.00 (from 2.3–3.5), reduces GC-target deviation by ∼82 % relative to CAI maximisation (4.0 vs 21.8 pp), and improves codon harmony by ∼83 % (0.77 vs 0.42), while shaping the translation-dynamics trajectory more closely to the biological ideal — at a deliberate, moderate cost in raw CAI (0.79 vs 1.00). It is the only strategy tested that holds all of these axes within acceptable bounds simultaneously. The platform’s conservative cassette-design philosophy — selecting from validated part libraries rather than inventing regulatory sequences — positions PDD as a scientifically defensible design aid, though all results are computational and wet-lab validation remains outstanding. All source code, benchmark data, and documentation are freely available.

## Funding

This work received no specific grant from any funding agency in the public, commercial, or not-for-profit sectors; it was supported by Voxelta Private Limited.

## Conflict of Interest

The authors declare no conflict of interest.

## Data and Code Availability

All data underlying this article are openly available under the MIT licence in the project repository. This includes the full source code, the reproducible benchmark suite, the committed Kazusa CUTG source data and its table generator, the figure-generation and PDF-build scripts, and the 172-test suite. Every figure, table, and benchmark number in this article can be regenerated from the repository with a fixed random seed; no proprietary data or external services are required. The repository is available at: https://github.com/Dinesh431786/Plant-DNA-Designer

## Author Contributions

Dinesh K conceived the project, designed the algorithms, implemented the software, and wrote the manuscript. H. Swetha contributed to algorithm design, literature review, and manuscript editing. Both authors read and approved the final manuscript.

## Notes

https://github.com/Dinesh431786/Plant-DNA-Designer

### Competing Interest Statement

The authors have declared no competing interest.

